# The expression of PKM1 and PKM2 in developing, benign, and cancerous prostatic tissues

**DOI:** 10.1101/2023.09.27.559832

**Authors:** Lin Li, Siyuan Cheng, Yunshin Yeh, Yingli Shi, Nikayla Henderson, David Price, Xin Gu, Xiuping Yu

## Abstract

Neuroendocrine prostate cancer (NEPCa) is the most aggressive type of prostate cancer. However, energy metabolism, one of the hallmarks of cancer, in NEPCa has not been well studied. Pyruvate kinase M (PKM), which catalyzes the final step of glycolysis, has two main splicing isoforms, PKM1 and PKM2. PKM2 is known to be upregulated in various cancers, including prostate adenocarcinoma (AdPCa). In this study, we used immunohistochemistry, immunofluorescence staining, and bioinformatic analysis to examine the expression of PKM1 and PKM2 in mouse and human prostatic tissues, including developing, benign and cancerous prostate. We found that PKM2 was the predominant isoform expressed throughout prostate development and PCa progression, with slightly reduced expression in some NEPCa samples.

PKM1 was mostly expressed in stromal cells but low-level PKM1 was also detected in prostate basal epithelial cells. Its expression was absent in the majority of PCa specimens but present in a subset of NEPCa. Additionally, we evaluated the mRNA levels of ten PKM isoforms that express exon 9 (PKM1-like) or exon 10 (PKM2-like). Some of these isoforms showed notable expression levels in PCa cell lines and human PCa specimens. These findings lay the groundwork for understanding PKMs’ role in PCa carcinogenesis and NEPCa progression. The distinct expression pattern of PKM isoforms in different PCa subtypes may offer insights into potential therapeutic strategies for treating PCa.

## 1. Introduction

The prostate is a male reproductive accessory gland. It comprises multiple cell types including luminal epithelial, basal epithelial and stromal cells^1^. Prostate cancer (PCa) primarily originates from luminal epithelial cells; thus the majority of PCa cases are prostate adenocarcinoma (AdPCa). Androgen deprivation therapy (ADT) is the first-line therapy for advanced AdPCa^2^. However, ADT eventually fails and PCa relapses and progresses into more aggressive castrate-resistant PCa^2^. With heightened androgen blockade, the more aggressive form, neuroendocrine prostate cancer (NEPCa), emerges^3^.

A hallmark of cancer is altered energy metabolism, often characterized by a preference for glycolysis, even in the presence of oxygen, which is known as the ‘Warburg effect’^4^. The Warburg effect signifies increased glycolytic flux and lactate production in cancer cells, aiding cancer progression^4^.

Glucose metabolism in the prostate, however, differs from other tissues. Normal prostate luminal epithelial cells consume a substantial amount of glucose to produce citrate for secretion, a specialized function of prostate epithelia^5^. At the molecular level, these cells accumulate zinc, which inhibits m-aconitase, an enzyme that converts citrate to isocitrate. The inhibition of m-aconitase minimizes citrate oxidation, leading to citrate secretion and a truncated tricarboxylic acid (TCA) cycle in normal prostates^5^.

In transformed PCa cells, however, glucose metabolism undergoes significant changes^5^. In these cells, zinc levels are reduced, which enables aconitase to catalyze the conversion of citrate to isocitrate^5^. Consequently, the metabolism shifts towards citrate oxidation, allowing the TCA cycle to proceed^5^. This alteration results in an increase of TCA cycle flux in PCa cells compared to normal prostate epithelial cells^5^. Because of this, the Warburg effect is only modest in early-stage PCa^5^. However, advanced AdPCa tumors, like many other solid tumors, exhibit more pronounced Warburg effect, which maintains high levels of glycolytic intermediates to support the biosynthesis required for the rapid proliferation of the malignant cells^5^. As for how glucose metabolism changes in NEPCa, it remains largely unknown.

A key enzyme that regulates glucose metabolism in cancer cells is pyruvate kinase muscle isozyme (PKM)^6,7^. PKM catalyzes the conversion of phosphoenolpyruvate (PEP) to pyruvate, one of the three irreversible and heavily regulated steps in glycolysis^6,7^. PKM has PKM1 and PKM2 isoforms, due to alternative splicing^6,7^. PKM1 includes exon 9 but not exon 10, whereas PKM2 includes exon 10 but not exon 9. PKM1 is predominantly expressed in energy-consuming tissues, such as heart, muscle, and brain, where it forms a constitutively active tetramer to promote oxidative phosphorylation^8,9^. In contrast, PKM2 expression is more prevalent in embryonic tissues and cancer cells^10–12^. Unlike PKM1, PKM2 can exist as a catalytically active tetramer or a less active dimer/monomer^7,13,14^, and its stoichiometry is regulated by allosteric factors and post-translational modifications^14,15^. PKM2 is highly expressed in various cancer types, including AdPCa, and contributes to increased glycolytic flux and tumorigenesis in cancer cells^6,7^. In these cells, PKM2 is predominantly expressed as dimer, which has lower pyruvate kinase activity, leading to the accumulation of glycolytic intermediates^6,7^.

Apart from PKM1 and PKM2, pyruvate kinase has two other isoforms, PKL and PKR, encoded by the PKLR gene under the control of different promoters^16,17^. PKL is mainly expressed in the kidney, liver, and intestine, while PKR is expressed predominantly in red blood cells^17^.

A recent study found that in prostate, PKM1 is mostly expressed in stromal cells, whereas PKM2 is expressed in both normal epithelia and cancer cells^18^. It has also been shown that the expression level of PKM2 is higher in prostate adenocarcinomas compared to normal human prostatic tissues and even higher in Gleason 8-10 tumors compared to Gleason 6-7 adenocarcinomas^19,20^. However, the expression of PKM1 and PKM2 in NEPCa, the most aggressive type of prostate cancer, has not been well studied. In this study, using immunohistochemistry (IHC) and immunofluorescence (IF) staining, we assessed the expression of PKM1 and PKM2 during prostate development, in PCa mouse models and human prostatic specimens including benign human prostate (BPH), prostate adenocarcinoma (AdPCa), and neuroendocrine prostate cancer (NEPCa).

## 2. Materials and Methods

### 2.1 Sample collection

Cell-derived xenograft tumors and prostatic tissues derived from wild type and TRAMP mice were from our archival collection. De-identified human prostate tissue specimens were obtained from LSU Health-Shreveport Biorepository Core, Overton Brooks VA Medical Center and Tissue for Research organization in accordance with LSU Health-Shreveport IRB protocols.

All methods were carried out in accordance with relevant guidelines and regulations. All experimental protocols were approved by LSU Health-Shreveport IRB. Informed consent was obtained from all subjects.

### 2.2 Immunohistochemistry and immunofluorescence staining

IHC staining was performed using Vectastain elite ABC peroxidase kit (Vector Laboratories, Burlingame, CA) as described previously^21^. Primary antibodies include PKM1 and PKM2 (7067S, dilution 1:500, and 4053S, dilution 1:1000, respectively, Cell Signaling Technology, Danvers, MA), p63 and FOXA2 (ab735 and ab108422, respectively, dilution 1:500, Abcam, Cambridge, MA), HOXB13 and chromogranin A (CHGA) (sc-28333 and sc-1488, respectively, dilution 1:200, Santa Cruz Biotechnology, Dallas, TX), FOXA1 (A15278, dilution 1:500, Abclonal, Woburn, MA), Synaptophysin (SYP) (611880, dilution 1:1000, BD biosciences, San Jose, CA), and NKX3.1 (0314, dilution 1:500, Athena Enzyme Systems, Baltimore, MD). Images were taken using a Zeiss microscope (White Plains, NY). The intensity of expression was graded as 0 (negative), 1+ (low), 2+ (moderate), and 3+ (high). For IF staining, primary antibodies include CHGA (AB_1553436, dilution 1:50, Developmental Studies Hybridoma Bank, Iowa City, IA), PKM1, PKM2, p63, and SYP (source was the same as mentioned above, dilution 1:100). The IF staining was imaged with a Nikon fluorescence microscope (Melville, NY) as reported previously.

### 2.3 Cell culture

PCa cell lines (VCap, LNCaP, C4-2B, 22RV1, PC3, and DU145) were obtained from ATCC. These cells were cultured in RPMI 1640 supplemented with 10% FBS, and 1% penicillin-streptomycin. H660 cells were cultured in Prostalife media (LifeLine Cell Technology, Oceanside, CA). C2C12 cells were cultured in DMEM supplemented with 10% FBS, and 1% penicillin-streptomycin. Cells were maintained in an incubator at 37 °C and 5% CO2.

### 2.4 Western blot

Cells were lysed in Laemmli SDS sample buffer followed by SDS-PAGE and Western blotting. Primary antibodies are beta-actin (sc-47778, dilution 1:500, Santa Cruz Biotechnology, Dallas, TX), PKM1 and PKM2 (the same source as mentioned above, dilution 1:1000), and HRP-conjugated secondary antibody (Cell Signaling, Beverly, MA). Protein bands were visualized by using ProSignal® Dura ECL Reagent (Genesee Scientific, San Diego, CA) and Chemidoc™ Touch Imaging System (Bio-Rad).

### 2.5 Cross-linking reaction

Cells were washed with ice-cold PBS three times and treated with 1-5mM disuccinimidyl suberate (DSS, A39267, Thermo Scientific, Waltham, MA) for 30 min at room temperature. The reaction was stopped by adding the quenching solution (1M Tris, PH 7.5) to the final concentration of 20mM for 15 min. Then, cell lysates were used for WB as described above.

### 2.6 Bioinformatics analysis

RNAseq data in the CTPC collection were analyzed to assess the expression of PKM isoforms using STAR-Salmon based transcript level quantification method^22–25^. The raw Fastq files were downloaded from NCBI SRA database and dbGaP (phs000909.v1.p1^26^, phs000915.v2.p2^27^). The raw count and TPM data of TCGA data were acquired from GDC Data Portal. All accession numbers were listed in Table S1. NCBI human GRCh38 (release 106) and associated GFT files were acquired through “AWS iGenomes” (https://ewels.github.io/AWS-iGenomes/) and were used for alignment. FastQC (https://github.com/s-andrews/FastQC), Trim Galore (https://github.com/FelixKrueger/TrimGalore), STAR^22^, Salmon^23^ software were used for sequencing quality control, alignment and quantification in the Linux Ubuntu environment. The transcript expression was visualized in R (V 4.3.0) environment. The TPM values were first log2 transformed and quantile normalized using “preprocessCore” R package^28^. The data was visualized by boxplot using “ggplot2” R package^29^.

## 3. Results

### 3.1. The expression of Pkm1 and Pkm2 in murine prostate

We first examined the expression of Pkm1 and Pkm2 during murine prostate development, including embryonic prostate undergoing budding morphogenesis (urogenital sinus, UGS), postnatal 2– and 3-week developing prostate, and fully developed 8-week prostate. The prostatic buds in UGS were confirmed by the expression of Foxa2 & p63 (Figs. 1C & 1D). The epithelial cells in postnatal prostate were indicated by the expression of Hoxb13 (Figs. 1G, 1K & 1O), Foxa1 (Figs. 1H & 1L), and Nkx3.1 (Fig. 1P). We found that Pkm1 expression was essentially absent in the developing prostate (Figs. 1A, 1E & 1I) but present in the developed prostate, albeit at a low level (Fig. 1M). Moreover, Pkm1 was predominantly expressed in the stromal cells of the developed prostate (Fig. 2A). In contrast, Pkm2 was predominantly expressed in prostatic epithelial cells throughout prostate development (Figs. 1B, 1F, 1J, 1N & 2G).

**Figure 1.**
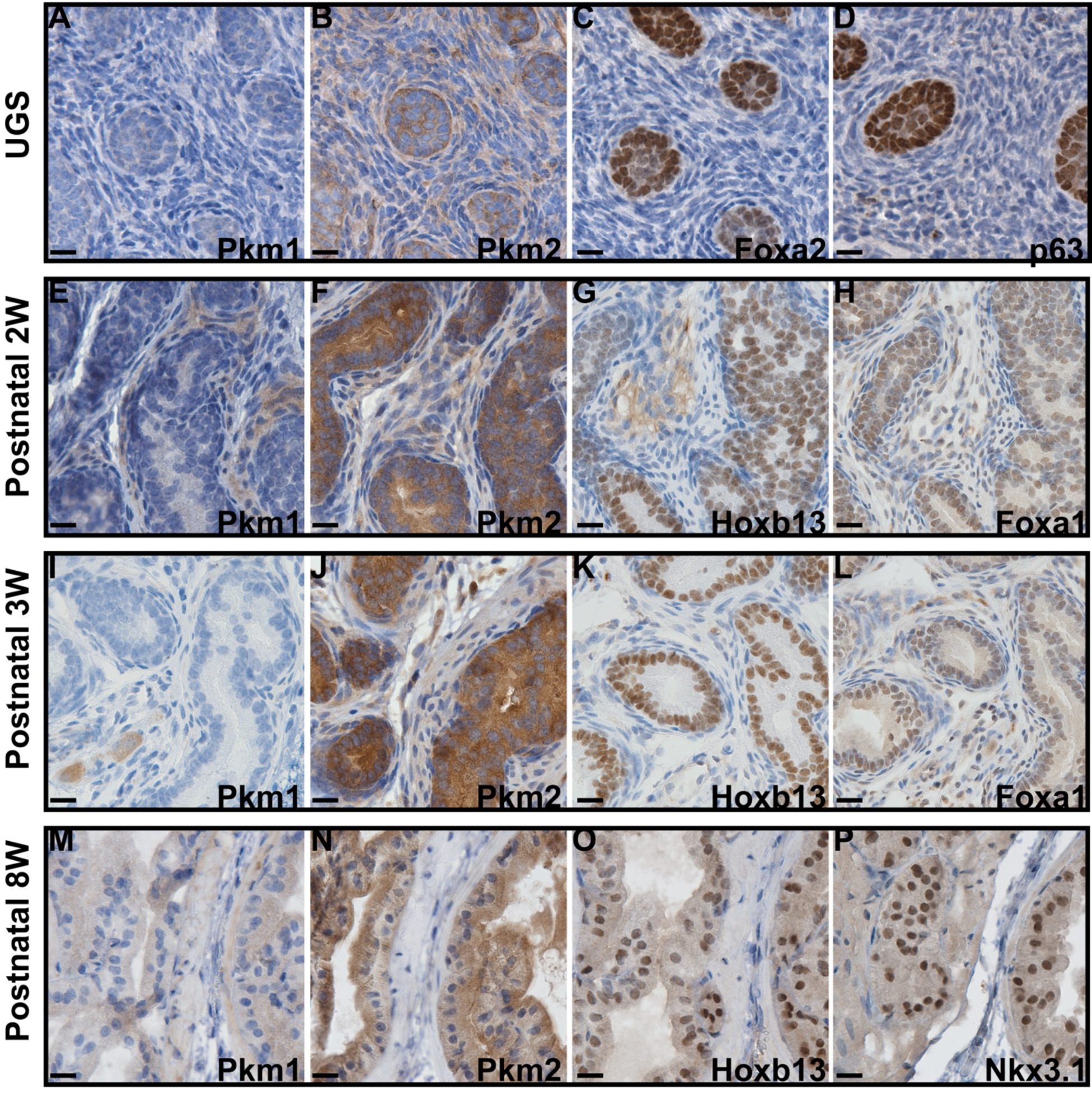
Immunohistochemistry staining to assess the expression of Pkm1 and Pkm2 in murine prostates at different developmental stages. The expression of Foxa2, p63, Foxa1, Hoxb13, and Nkx3-1 highlights the embryonic prostate buds and postnatal prostate luminal epithelial cells. Scale bar = 20 μM.

**Figure 2.**
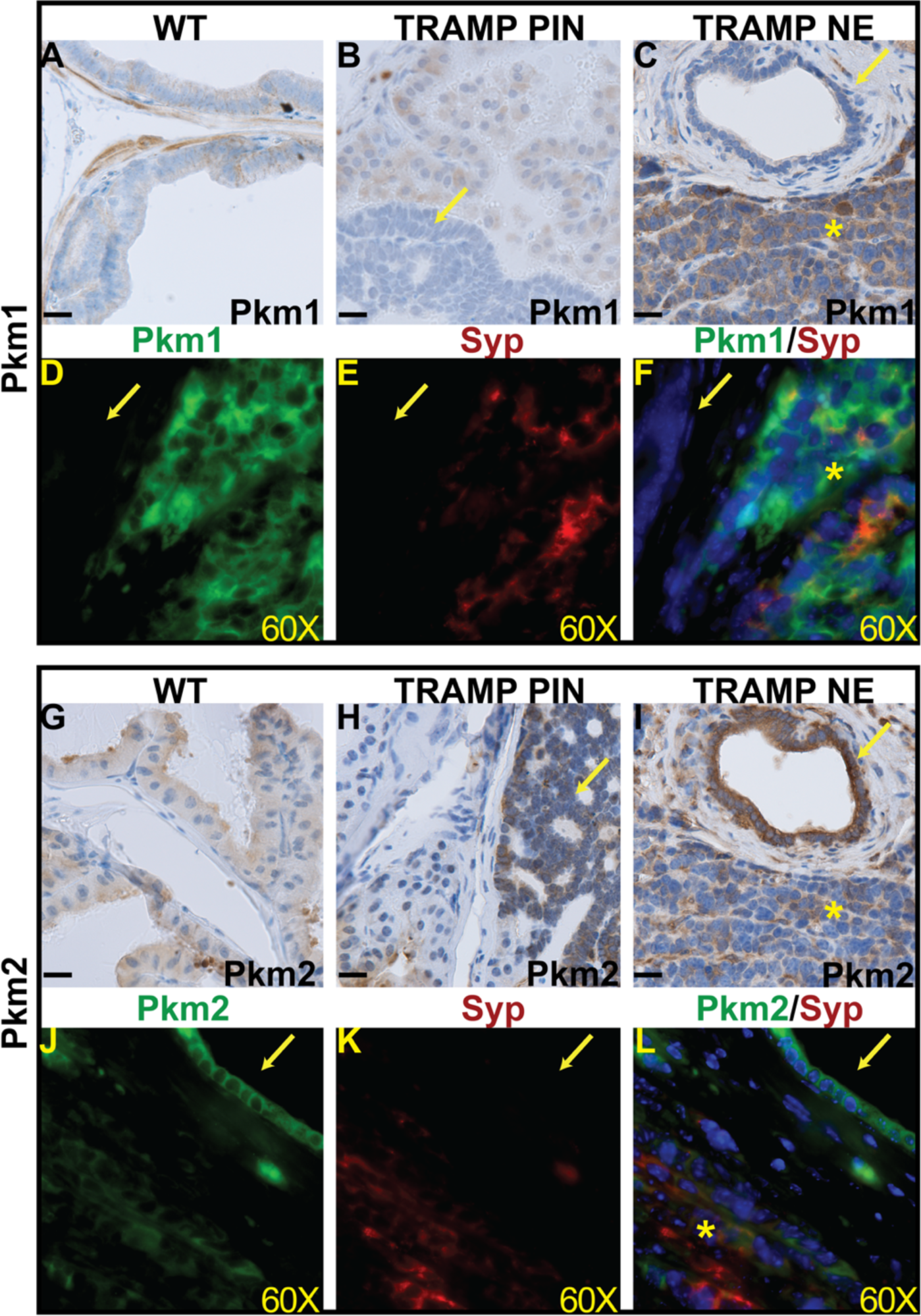
The expression of Pkm1 and Pkm2 in prostate cancer mouse models. (**A**-**C**) IHC staining of Pkm1 in wild-type prostate (**A**), TRAMP PIN (**B**) and TRAMP NEPCa tumors (**C**). (**D**-**F**) Dual IF staining of Pkm1 (**D**, green) with NE marker Syp (**E**, red)) in TRAMP tumor. Pkm1 is expressed in NEPCa area but not in adjacent PIN. (**G**-**I**) IHC staining of Pkm2 in wild-type (**G**), TRAMP PIN (**H**) and TRAMP NEPCa (**I**). (**J**-**L**) Dual IF staining of Pkm2 (**J**, green) with NE marker Syp (**K**, red)) in TRAMP tumors. Pkm2 level is relatively lower in NEPCa area compared with adjacent PIN. The yellow arrows denote prostatic intraepithelial neoplasia (PIN) lesions. Astrids denote NEPCa. Scale bars for IHC = 20 μM. IF images were taken using 60X lens.

Next, we assessed the expression of Pkm1 and Pkm2 during murine PCa progression using the transgenic adenocarcinoma of the mouse prostate (TRAMP), a widely used PCa mouse model^30^. TRAMP mice develop prostatic intraepithelial neoplasia (PIN), and a subset of tumors progress into NEPCa^31^. We found that Pkm1 expression was present in the epithelial cells of wild-type prostates (Fig. 2A) as well as normal adjacent cells in TRAMP tumors (Fig. 2B) but absent in the PIN lesions of TRAMP tumors (Fig. 2B). In the TRAMP tumors that have progressed to NEPCa, Pkm1 was expressed in the NEPCa areas while its expression remained absent in the PIN lesions (Fig. 2C). In contrast, high Pkm2 expression was detected in both the PIN lesions and NEPCa cells of TRAMP tumors (Figs. 2H & 2I). It is noteworthy that the expression level of Pkm2 was higher in PIN lesions than the adjacent NEPCa areas in TRAMP tumors (Fig. 2I). To confirm the differential expression of Pkm1 and Pkm2 in TRAMP tumors, we performed dual immunofluorescence staining of Pkms with NEPCa marker Synaptophysin (Syp). Pkm1 expression was detected in the NEPCa cells (indicated by Syp positivity) but not in the adjacent PIN lesion (Figs. 2D-2F). In contrast, Pkm2 expression was more prominent in the PIN lesions than the adjacent NEPCa areas (Figs. 2J-2L).

### 3.2. PKM1 expression is absent in most AdPCa but present in a subset of NEPCa, while PKM2 expression is abundant in most human prostatic tissues

We also conducted immunohistochemical staining to assess the expression of PKM1 and PKM2 in human prostatic tissues, including benign prostatic hyperplasia (BPH), low-grade AdPCa, high-grade AdPCa, as well as NEPCa. Consistent with Pkms’ expression pattern in murine prostate, PKM1 was predominantly expressed in stromal cells (Figs. 3A & 3E). In BPH, PKM1 expression was detected in basal epithelial cells as well (Fig. 3A, n=13). This expression pattern was confirmed by dual immunofluorescence staining of PKM1 with p63, a marker of prostate basal epithelia. The result showed a high expression of PKM1 in stromal cells, with detectable expression in basal epithelial cells (Figs. 3E-H). In low-grade and high-grade human prostate adenocarcinoma (AdPCa), PKM1 expression was generally scarce (Figs. 3B & 3C, n=12 and 14, respectively). However, in the high-grade AdPCa cases with NE differentiation, PKM1 was expressed in the rare NE cells, which were indicated by the co-expression of NEPCa marker Chromogranin A (CHGA, Figs. 3I-L). Additionally, PKM1 expression was detected in 5/12 human NEPCa samples (Fig. 3D & 3Q). The expression of PKM1 in PCa epithelial cells, including both AdPCa and NEPCa, was quantified and its distribution among PCa samples is presented in Fig. 3Q.

**Figure 3.**
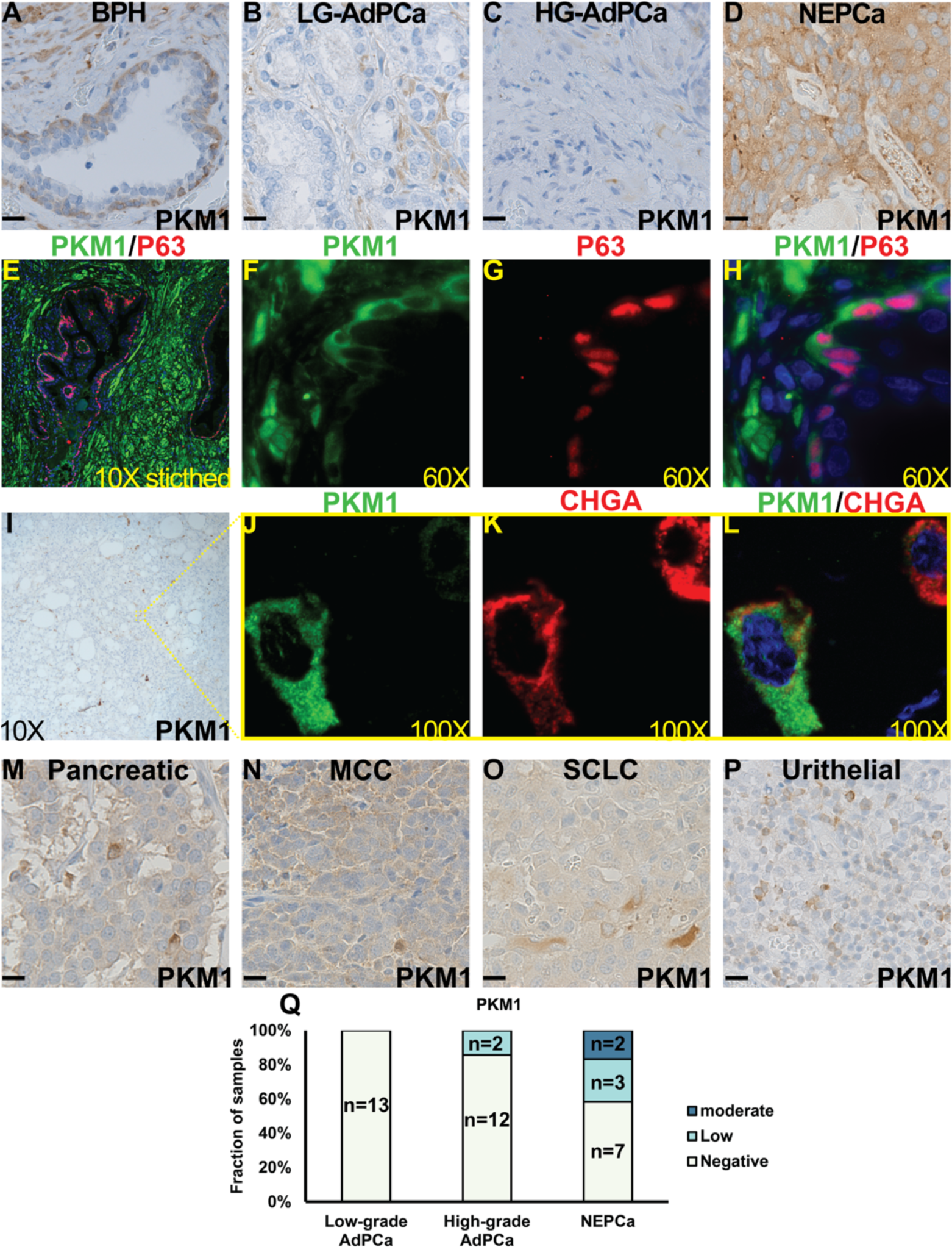
The expression of PKM1 in human prostate specimens and other types of NE tumors. (**A-D**) IHC staining of PKM1 in BPH (**A**), low-grade (LG) AdPCa (**B**), high-grade (HG) AdPCa **(C)** and NEPCa **(D**). The quantification of PKM1 IHC staining intensity in human PCa is shown in (**Q**). (**E-H**) IF staining of PKM1 (**F**, green) and basal epithelial cell marker p63 (**G**, red) in human BPH. PKM1 is expressed in prostate basal epithelial and stromal cells. (**I-L**) Serial sections derived from an AdPCa with NED were used for IHC staining of PKM1 (**I**) or dual IF staining of PKM1 (**J**, green) and NE marker CHGA (**K**, red). PKM1 is co-expressed with CHGA in some scattered NEPCa cells. (**M-P**) IHC staining of PKM1 in pancreatic adenocarcinoma with NED (**M**, n=4), Merkel cell carcinoma (MCC) (**N**, n=3), small cell lung cancer (**O**, n=2), and urothelial NED samples (**P**, n=3). Scale bar for IHC = 20 μM.

Moreover, we examined the expression of PKM1 in other types of NE tumors, including pancreatic adenocarcinoma with NE differentiation (Fig. 3M, 4/4 positive), Merkel cell carcinoma (Fig. 3N, 2/3 positive), small cell lung cancer (SCLC, Fig. 3O, 1/2 positive), and urothelial NE carcinoma (Fig. 3P, 2/3 positive). We found that PKM1 was expressed in a subset of these NE tumors as well, highlighting its potential role in the pathogenesis of neuroendocrine tumors.

On the other hand, the expression of PKM2 was detected in the majority of human prostatic tissues. In the BPH samples, PKM2 expression was prominent in basal epithelial cells and comparatively moderate in luminal epithelial cells (Figs. 4A & 4B, n=14). This expression pattern was confirmed by dual immunofluorescence staining, which demonstrated that PKM2 was highly expressed in basal epithelial cells (indicated by the expression of basal marker p63) with less staining in luminal epithelial cells (Figs. 4I-L). Human AdPCa samples exhibited a diverse PKM2 expression pattern, varying from negative to high levels in both low-grade (Figs. 4C& 4D, n=12) and high-grade AdPCa (Figs. 4E & 4F, n=15). Additionally, PKM2 was expressed in all the human NEPCa samples examined (Figs. 4G & 4H, n=9). The intensity of PKM2 staining in PCa cells, including both AdPCa and NEPCa, was quantified and its distribution among PCa samples is presented in Fig. 4M.

**Figure 4.**
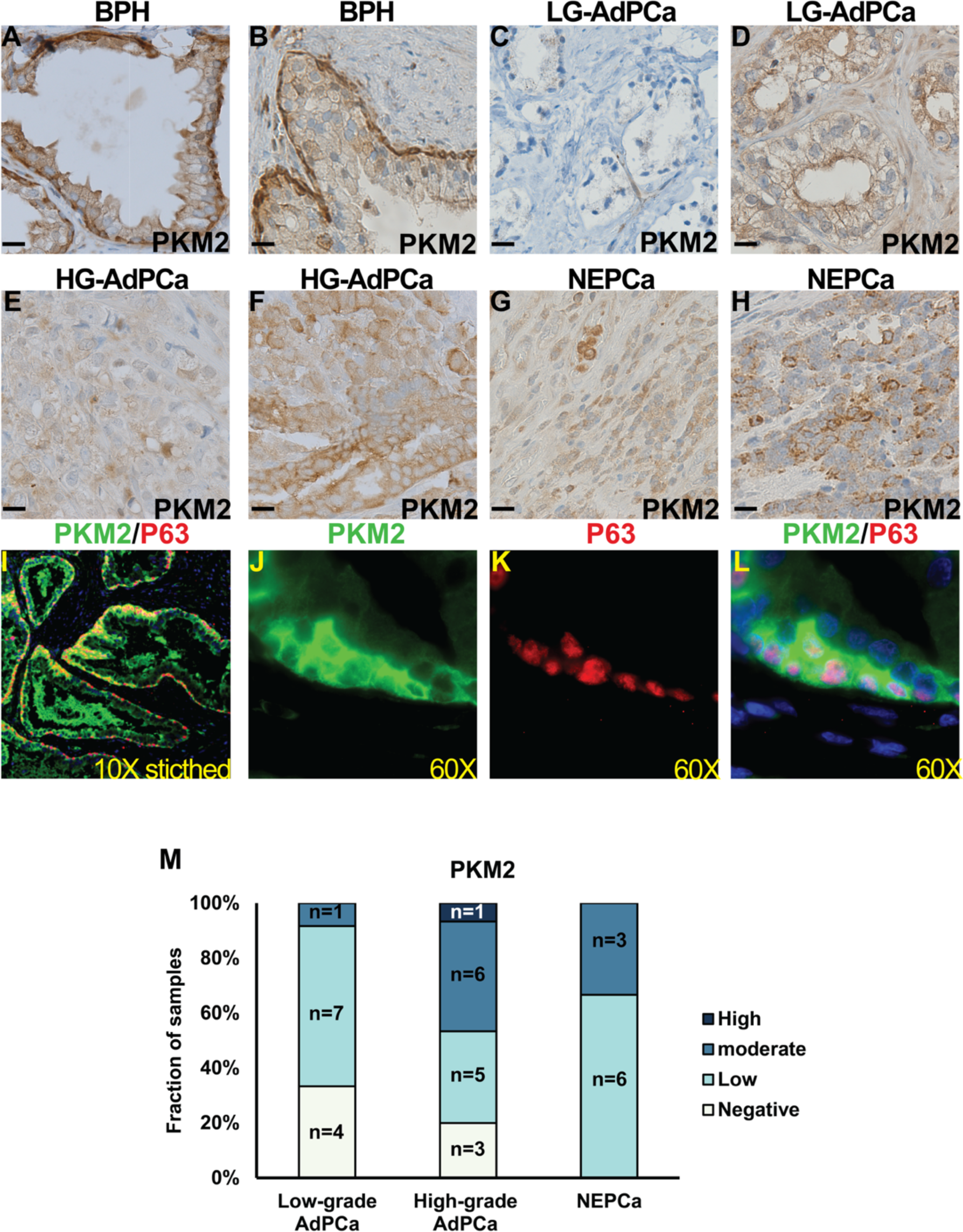
The expression of PKM2 in human prostate specimens. (**A-H**) IHC staining of PKM2 in BPH (**A** & **B**), low-grade (LG) AdPCa (**C** & **D**), high-grade (HG) AdPCa (**E** & **F**) and NEPCa (**G** & **H**). **(I-L)** IF staining of PKM2 (**B**, green) and basal epithelial cell marker p63 (**C**, red) in human BPH. (**M**) Quantification of PKM2 staining intensity. Scale bar for IHC = 20 μM.

Taken together, PKM2 was predominantly expressed in human prostatic tissues, ranging from benign to cancerous prostate, including both AdPCa and NEPCa. PKM1 expression in PCa was limited but notable, showing low to moderate expression in a subset of NEPCa cases.

### 3.3. The expression of PKM1 and PKM2 in human PCa cell lines

Using IHC staining and Western blot analysis, we evaluated the expression of PKM1 and PKM2 in human PCa cell lines, including AR positive (VCaP, LNCaP, C42B and 22RV1), AR negative (PC3 and DU145), and NEPCa (H660) cell lines. Both cultured cells and cell-derived xenograft tumors were used. Our IHC staining demonstrated that the expression of PKM1 was lowest in 22RV1– and highest in PC3-derived xenograft tumors (Figs. 5A-G). This expression pattern was consistent with the Western blot analysis of the lysates derived from *in vitro*-cultured cells (Fig. 5H). NEPCa H660 cells expressed moderate level of PKM1 (Fig. 5H). PKM2 expression was consistently detected in all seven cell lines (Figs. 5I-P).

**Figure 5.**
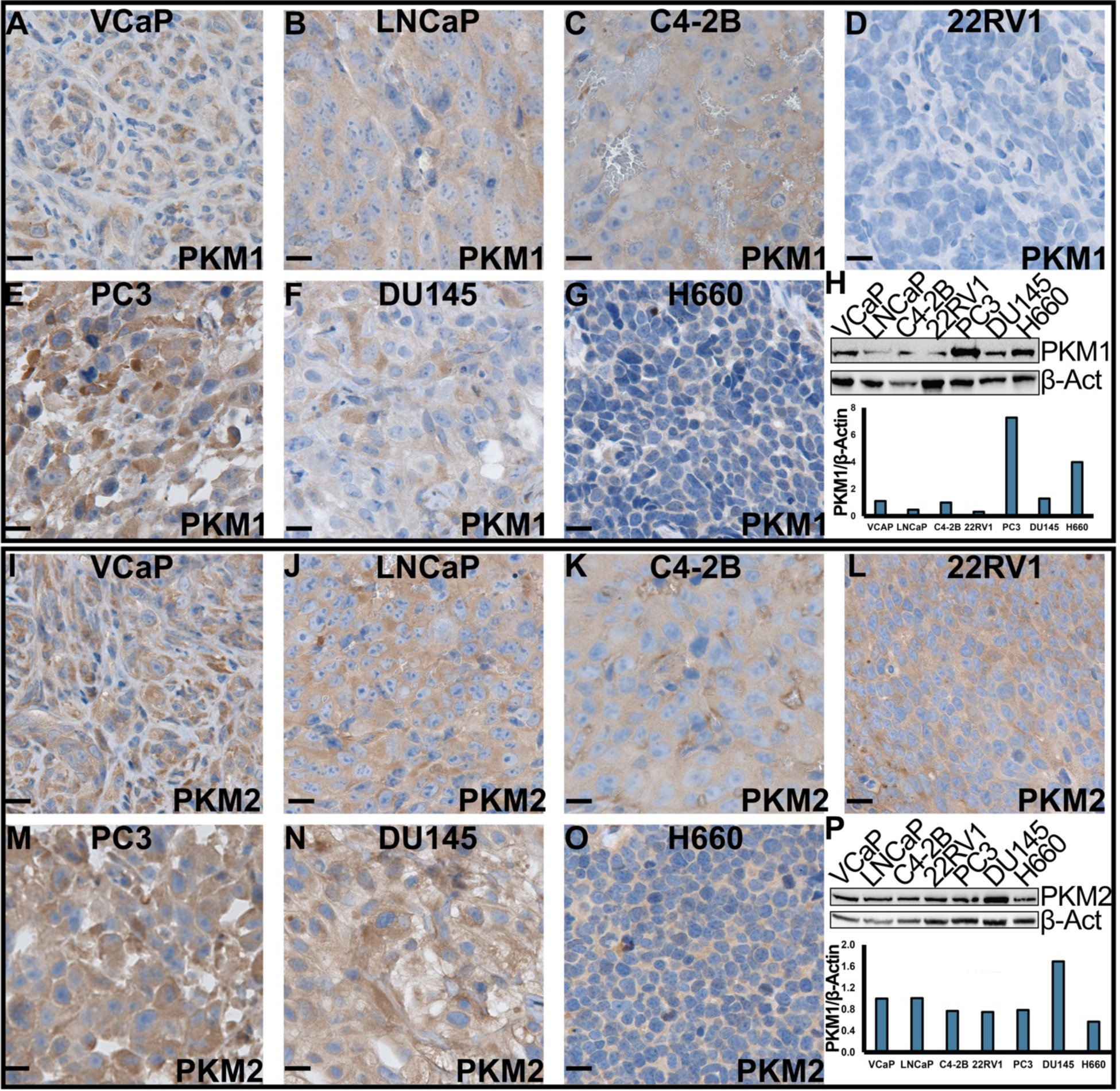
The expression of PKM1 and PKM2 in human PCa cell lines. (**A-G** and **I-O**) IHC staining of PKM1 and PKM2, respectively, in xenograft tumors derived from human PCa cell lines. Scale bar = 20 μM. (**H** & **P**) Western blotting analysis of PKM1 (**H**) and PKM2 (**P**) in human PCa cell lines.

### 3.4 Bioinformatic analysis reveals the expression of PKM1/2 transcripts in PCa cell lines and patient samples

The PKM1/2 transcripts are classified into PKM1 and PKM2 based on the alternative splicing of a single exon, which is resulted from an exclusive skipping of exon 9 or 10. Antibodies are available to distinctively target exon 9 or exon 10, but these antibodies cannot differentiate the various isoforms of each PKM class. The development of high throughput RNA sequencing and associated bioinformatic tools enables analyzing each of these isoforms. Using publicly available RNAseq data generated from human PCa cell lines and patient samples, we quantified the transcripts of 10 PKM isoforms including the previously well-studied PKM1 (NM_182470.2 and NM_182471.2) and PKM2 (NM_002654.4) as well as the under studied isoforms.

The ten PKM isoforms were presented in Fig. 6A, and the expression levels of each isoform in human prostatic tissue specimens were summarized in Fig. S1. Based on the inclusion of exon 9 or exon 10, we classified these isoforms into “PKM1-like” and “PKM2-like” transcripts, respectively. The PKM1 and PKM2 antibodies used in this study target exon 9 or 10; thus, they detect the protein products of PKM1-like or PKM2-like transcripts. The PKM1-like transcripts consist of 4 isoforms, among which the NM_182470.2 and NM_182471.2 transcripts encode the same protein since the only difference in them lies within their 5’ UTRs. The protein encoded by these two isoforms is known as PKM1. There are six PKM2-like isoforms, among which the NM_002654.4 transcript encodes a protein that is known as PKM2. To coordinate our bioinformatics analysis of the ten PKM isoforms with the well-known PKM1 and PKM2, we summed the normalized log2-transformed TPM values of NM_182470.2 and NM_182471.2 as PKM1 transcript. The expression data of NM_002654.4 was directly used to reflect the levels of PKM2 transcript. Additionally, to complement our bioinformatics analysis with the PKM1/2 antibodies’ staining results, we summed the values of all exon 9-retained transcripts as PKM1-like expression data and all exon 10-retained transcripts as PKM2-like expression data. Overall, the levels of PKM2-like transcripts were higher than that of PKM1-like transcripts in both PCa cell lines and human PCa specimens (Figs. 6B & 6C). The NM_002654.4 transcript (PKM2) was the dominant PKM isoform expressed in these tissues (Fig. S1).

**Figure 6.**
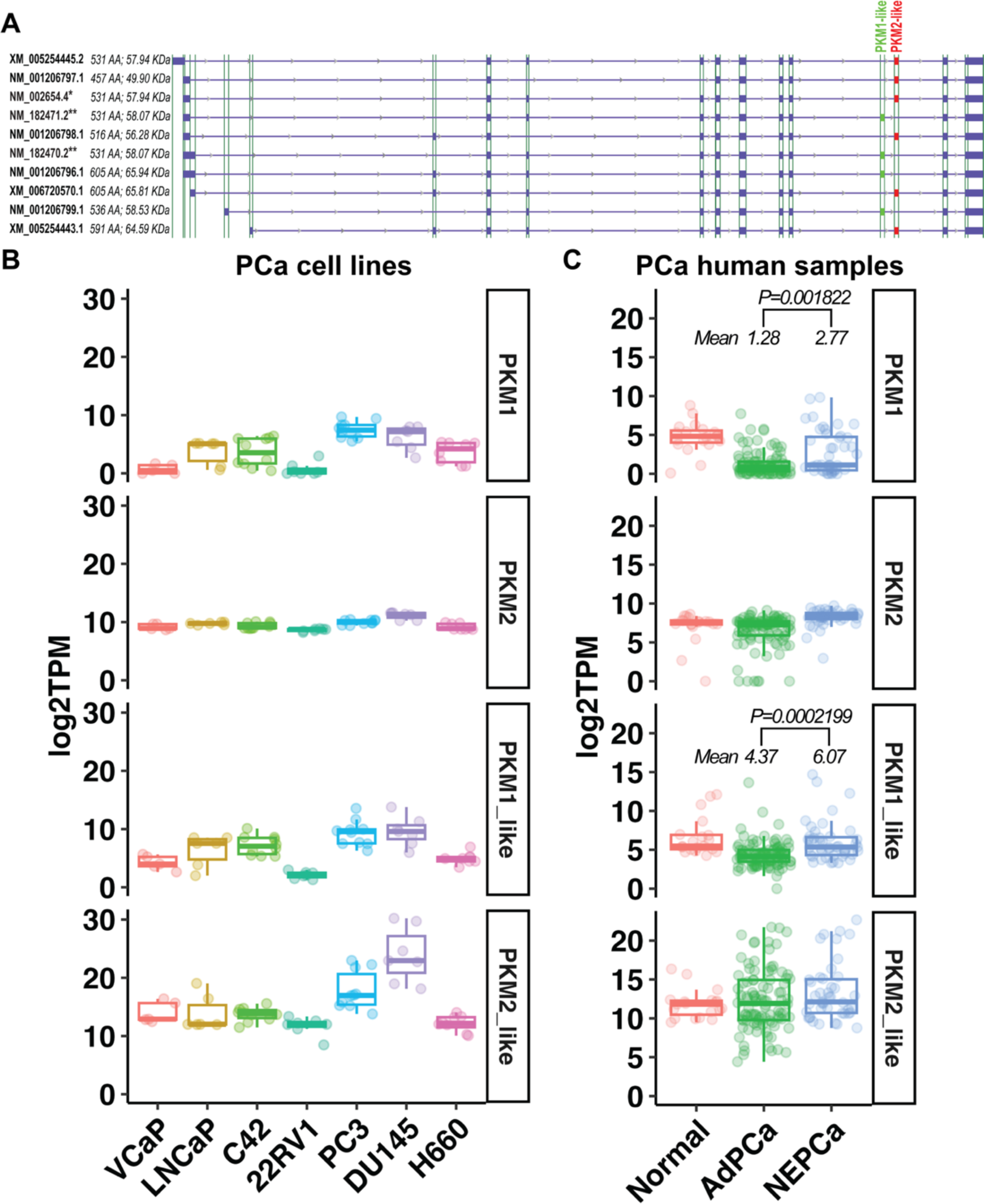
The expression of PKM1-like and PKM2-like isoforms in human PCa. (**A**) Ten PKM isoforms. PKM1-like isoforms are labeled in green and PKM2-like isoforms are in red. ** indicates the previously known PKM1 isoforms, and * indicates the previously known PKM2 isoform. (**B** & **C**) The levels of the previously known PKM1 and PKM2 as well as the newly classified PKM1-like and PKM2-like transcripts in human PCa cell line and human prostatic tissues.

Additionally, we analyzed the prostatic expression of PKL/R, the other two PK isoforms^25^. We found that the levels of PKL/R transcripts were generally low in human PCa cell lines and human PCa samples, but these transcripts were detected in a subset of human NEPCa (Fig. S2). This is consistent with a recent study that shows PKL/R expression is increased in castrate-resistant PCa and the upregulated PKL/R, coupled with elevated MYCN, may drive metabolic reprogramming during NE differentiation^32^.

### 3.5 The stoichiometry of PKM2

PKM1 forms constitutively active tetramers^33^. In contrast, PKM2 can exist as monomer, dimer, or tetramer. To study the stoichiometry of PKM2 in PCa, we conducted a cross-linking reaction. C2C12 myoblast cells were used as a positive control for the detection of PKM2 tetramers (Fig. 7). We found that in PCa cells, PKM2 formed inactive monomers/dimers but not active tetramers (Fig. 7).

**Figure 7.**
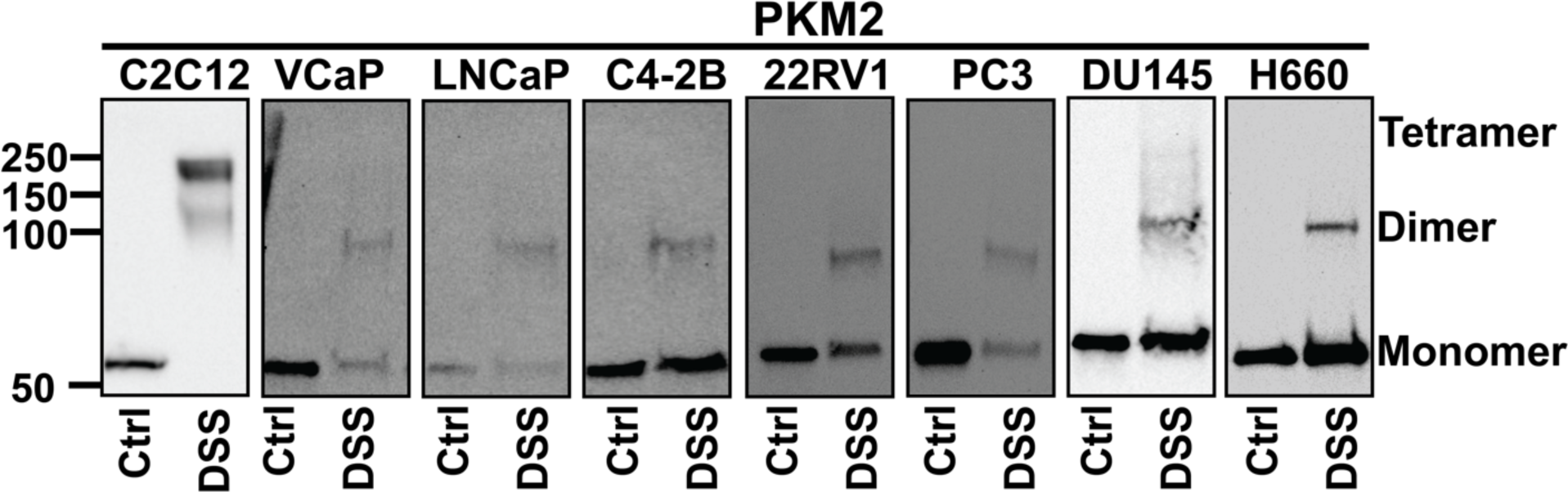
The stoichiometry of PKM2. Cells were treated with DSS reagent to form intramolecular crosslinks and Western blot analysis was conducted to examine the subunit association of PKM2. PKM2 forms monomers and dimers but not tetramers in PCa cells. C2C12 cells were used as a positive control for the detection of PKM2 tetramers.

## 4. Discussion

There is a paucity of NEPCa specimens because surgery is not the primary treatment option for late-stage PCa patients. This has limited research that can be pursued on energy metabolism in NEPCa. In this study, using the human prostatic samples we have collected as well as archived murine PCa tissues, we assessed the expression of PKM isoforms during prostate development and PCa progression. Our results show that PKM2 is the predominant isoform expressed in prostatic tissues and PKM1 expression is less prominent in prostate epithelia or AdPCa cells. Consistent with previous reports^18,34^, PKM1 is highly expressed in the stromal cells in both benign and cancerous prostatic tissues. Prostatic basal epithelial cells express ample PKM1 as well, suggesting these cells may have distinct glucose metabolism. Basal cells are absent in PCa, and PKM1 expression is restricted to the stromal regions in the tumors. This expression pattern suggests PKM1 may play an insignificant role in glucose metabolism in AdPCa.

In contrast, PKM2 is expressed in prostate epithelial cells. Its level is generally higher in high-grade AdPCa compared with normal prostate or low-grade AdPCa. Given the well-established function of PKM2 in regulating glucose metabolism, elevated PKM2 in high-grade AdPCa aligns well with the increased Warburg effect in these tumors. This is corroborated by the stoichiometry analysis of PKM2. While PKM1 forms constitutively active tetramer, PKM2’s stoichiometry is regulated by its allosteric regulators, such as serine and FBP, as well as post-translational modifications, such as phosphorylation and oxidation^7,13,14^. By crosslinking the proteins followed by Western blot analysis, we found that PKM2 forms less-active monomers and dimmers in PCa cells. This indicates that PKM2, the predominant isoform of PKM in PCa cells, exists in the stoichiometry of low pyruvate kinase activity. This could lead to the accumulation of glycolysis intermediates and the production of lactate in cancer cells^39,41,42^. This notion is consistent with previous studies that have shown PKM2 knockdown reduces lactate production and inhibits cell proliferation in DU145 cells and that Pkm2 knockout suppresses tumor progression in Pten null mice^18,20^. Taken together, these studies indicate that PKM2 promotes the Warburg effect and carcinogenesis in AdPCa.

Importantly, we found that PKM1 is co-expressed with NEPCa markers in human PCa tissues that have NE differentiation, but its expression is absent in the adjacent non-NEPCa cells. Additionally, PKM1 is expressed in a subset of NEPCa tumors, albeit at a low level. Moreover, we found that PKM1 is expressed in other types of NE tumors as well, including small cell lung cancer. This is consistent with literatures that report PKM1 expression in pulmonary NE tumors^35^. In contrast, PKM2 levels are notably lower in NEPCa than AdPCa. These results suggest that the glucose metabolism in NEPCa may differ from that in AdPCa and PKM1 may play a role in the energy metabolism in NEPCa.

Using publicly available RNAseq data, we analyzed the expression of ten PKM isoforms. We found that the prevalent isoforms expressed in prostatic tissues are PKM1 (NM_182470.2 & NM_182471.2) and PKM2 (NM_002654.4) transcripts. The antibodies that specifically target exon 9 or 10 can differentiate the PKM1-like from PKM2-like transcripts but not the individual isoforms. It is noteworthy that the PKM isoforms encode proteins of different molecular weight, which may enable the detection of certain isoforms within the PKM1-like and PKM2-like groups by Western blotting. However, our Western blot results only displayed a single band of PKM1 or PKM2, suggesting that the prevailing protein isoforms in PCa are PKM1 encoded by the NM_182470.2 & NM_182471.2 transcripts and PKM2 encoded by the NM_002654.4 transcript.

In summary, our study characterized the expression of PKM1 and PKM2 in prostatic tissues including developing, benign, and cancerous prostate tissues. This information contributes to our understanding of the metabolic alterations in aggressive PCa and may potentially lead to the development of targeted therapies to combat this deadly disease. However, due to the scarcity of NEPCa specimens, further research is needed to fully comprehend the metabolic profile of this aggressive PCa subtype and explore potential therapeutic avenues.

## Acknowledgements

This research was supported by NIH R01 CA226285, LSU Collaborative Cancer Research Initiative, LSU Health Shreveport Office of Research, and LSU Health Shreveport FWCC Stimulus grants to X. Yu, LSU Health Shreveport FWCC Carroll Feist postdoctoral Fellowship to S. Cheng, and LSU Health Shreveport FWCC Carroll Feist predoctoral Fellowship to L. Li. RNA sequencing data were downloaded from dbGaP through dbGaP accession number phs000909 and phs000915^7,7^. The data were generated under the support of NHGRI grant # U54 HG003067 to E. Lander at Broad Institute and SU2C/PCF Prostate Dream Team Translational Cancer Research Grant.

## Data Availability

The datasets analyzed in this study are available in the GEO (https://www.ncbi.nlm.nih.gov/geo/), dbGaP (https://www.ncbi.nlm.nih.gov/gap/) and GDC Data Portal (https://portal.gdc.cancer.gov/). The detailed accession information was listed in Table S1.

**Figure S1.**
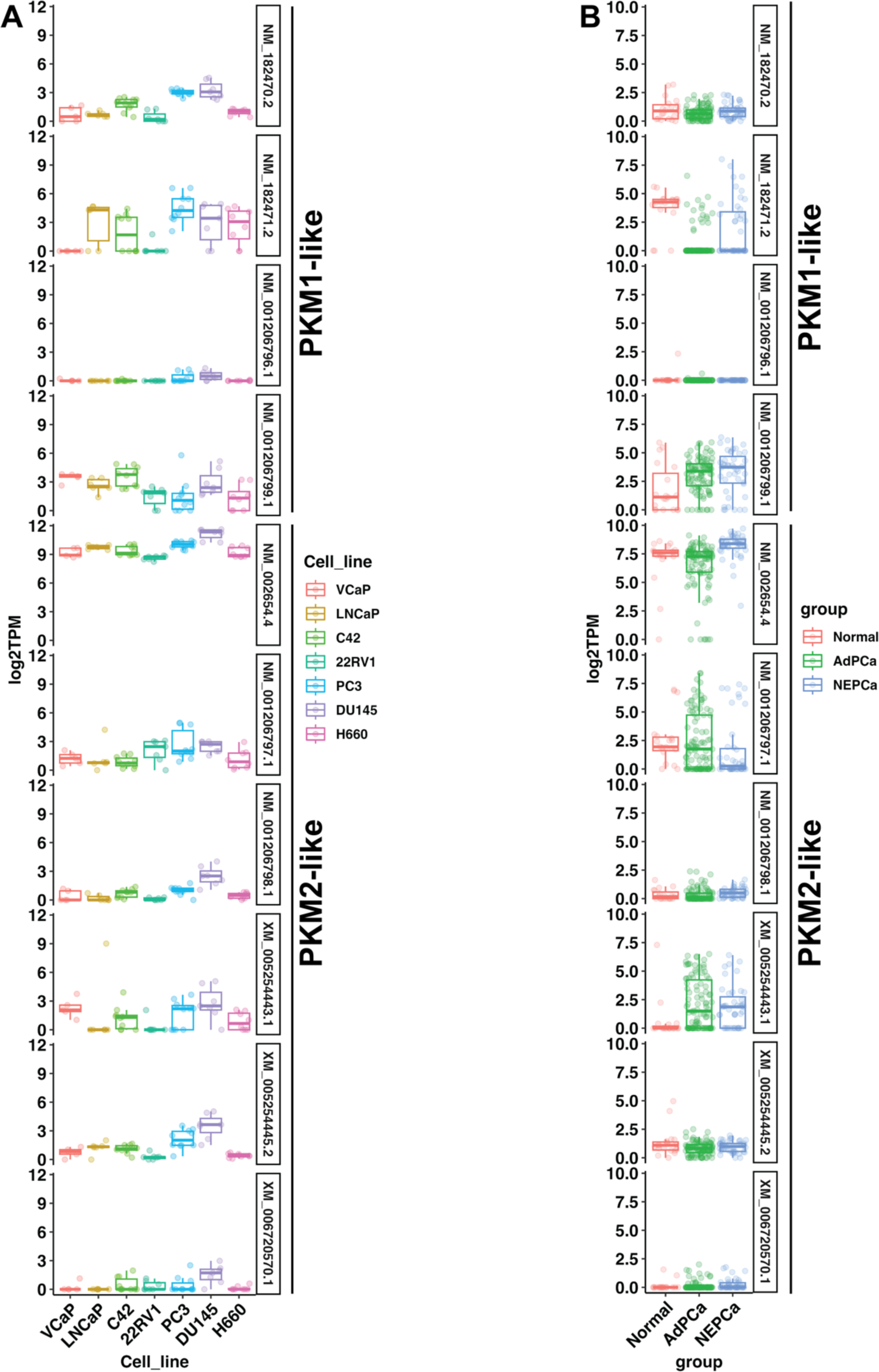
The mRNA expression of PKM isoforms in PCa cell lines (A) and human prostate Specimen (B).

**Figure S2.**
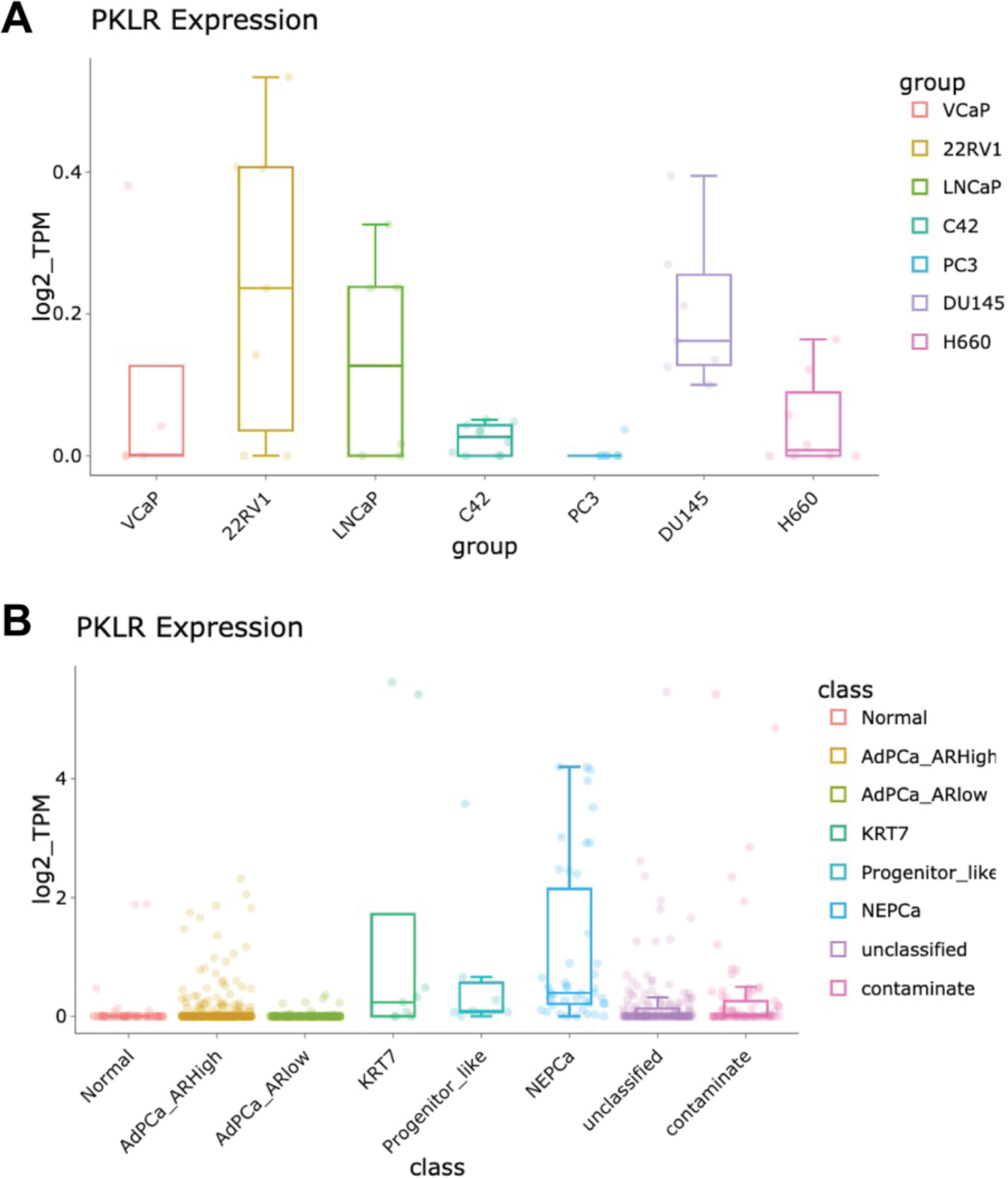
The mRNA expression of PKLR in PCa cell lines (A) and human prostate Specimen (B).

